## Supplementary information for "Cryo-EM structures of HKU2 and SADS-CoV spike glycoproteins and insights into coronavirus evolution"

### Materials and Methods

#### Expression and purification of HKU2 and SADS-CoV spike ectodomains

The cDNAs encoding HKU2 spike (YP_001552236) and SADS-CoV spike (AVM41569.1) were synthesized with codons optimized for insect cell expression. HKU2 ectodomains (1-1066) and SADS-CoV ectodomains (1-1068) were separately cloned into pFastBac-Dual vector (Invitrogen) with C-terminal foldon tag for trimerization and Strep tag for purification in the frame of PH promoter. They were expressed in Hi5 insect cell using Bac-to-Bac baculovirus system (Invitrogen) and purified as previously described[21,37]. Briefly, the construct was transformed into DH10Bac competent cells and the extracted bacmid was transfected into Sf9 cell using Cellfectin II reagent (Invitrogen). The baculoviruses were harvested after 7-9 days. The high-titer viruses were generated after one more amplification which used to infect Hi5 cells at a density of $1.5\times 10^6$/ml. After 60h in culture, the cell medium containing spike ectodomains were concentrated and exchanged to binding buffer (50mM Tris, pH 8.0, 150mM NaCl). The spike ectodomains were purified by StrepTactin (IBA) and then purified by gel-filtration chromatography using Superose 6 gel filtration column (GE Healthcare) pre-equilibrated with HBS buffer (10mM HEPES PH 7.2, 150mM NaCl). Purified ectodomains were concentrated for electron microscopy analysis.

#### Cryo-electron microscopy

Aliquots of spike ectodomains (4ul, 0.33mg/ml, in buffer containing 10mM HEPES PH 7.2, 150mM NaCl) was applied to glow-discharged holey carbon grids (Quantifoil grid, Au 300 mesh, R1.2/1.3). The grids were then blotted and then plunge-frozen in liquid ethane using FEI Vitrobot system (FEI).

Images were recorded using FEI Titan Krios microscope operating at 300 kV with a Gatan K2 Summit direct electron detector (Gatan Inc.) at Tsinghua University. The automated software (AutoEMation) was used to collect 7663 movies for HKU2 and 4568 movies for SADS-CoV at 130000 magnification and at a defocus range between 1-3 μm. Each movie has a total accumulate exposure of $49.784e^{-}/A^2$ fractionated in

32 frames of 175ms exposure. Data collection statistics are summarized in Supplementary Table 1.

Whole frames in each movie were corrected for beam-induced motion using MotionCo2[48]. The final image was bin averaged to give a pixel size of 1.061Å. The parameters of contrast transfer function (CTF) was estimated for each micrograph using GCTF[49]. Particles were automatically picked using Gautomatch (http://www.mrc-lmb.cam.ac.uk/kzhang/) and extracted using RELION[50]. Initially, ~1,400,000 particles for HKU2 and ~900,000 particles for SADS were subjected to 2D classification. After two or three additional 2D classification, the best class consisting ~750,000 particles (HKU2) and ~320,000 particles (SADS-CoV) were applied for creating 3D initial model, 3D refinement and 3D classification. A total of 421,490 particles (HKU2) and 152,334 particles (SADS-CoV) of best class were selected and subjected to 3D refinement with C3 symmetry to generate density map. The reported resolutions based on the gold-standard Fourier shell correlation (FSC) cutoff of 0.143 criterion were 2.38 Å for HKU2 spike and 2.83 Å for SADS-CoV spike after RELION post-processing. Local resolution variations were estimated using ResMap[51]. Data processing statistics are summarized in Supplementary Table 1.

**Model building and refinement**

As for the HKU2 spike model building, the initial model of HKU2 S1 NTD was generated using the SWISS-MODEL[52] and fit into the map using UCSF Chimera[53]. The other part of HKU2 spike ectodomains was obtained using Map to Model in PHENIX suit[54]. As for the SADS-CoV spike model building, the initial model was obtained by fit HKU2 S2, NTD, CTD separately into the map using UCSF Chimera[53]. Manual model rebuilding was carried out using Coot[55] and refined with PHENIX real-space refinement[54]. The quality of the final model was analyzed with Molprobity[56] and EMRinger[57]. The validation statistics of the structural models are summarized in Supplementary Table 1.

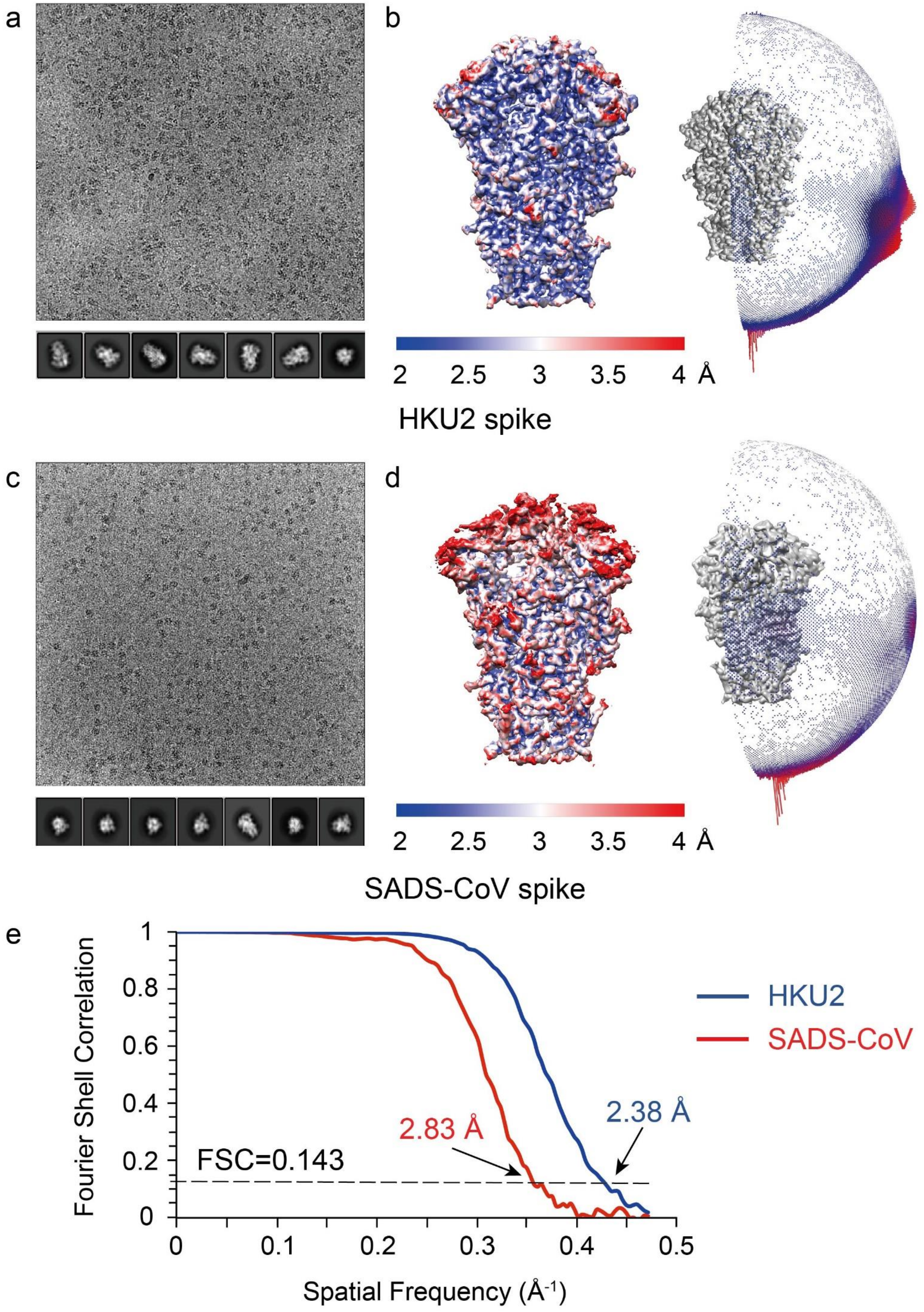


**Supplementary Fig. 1 Cryo-EM data and the statistics of the final density maps. (a)** Representative cryo-EM micrograph and selected 2D class averages of HKU2 spike. **(b)** Local resolution (Å) plotted on the cryo-EM map (left panel) as a heat map and particle orientation distribution (right panel) of HKU2 spike. The cryo-EM map is contoured at 4 RMS. The color scale from 2 Å to 4 Å is shown at the bottom of the local resolution map. The Red cylinders in the right panel represent the particles on these orientations; heights of cylinders represent the relative numbers of particles. **(c)** Representative cryo-EM micrograph and selected 2D class averages of SADS-CoV spike. **(d)** Local resolution (Å) plotted on the cryo-EM map (left panel) as a heat map and particle orientation distribution (right panel) of SADS-CoV spike. The cryo-EM map is contoured at 5.5 RMS. The color scale from 2 Å to 4 Å is shown at the bottom of the local resolution map. The Red cylinders in the right panel represent the particles on these orientations; heights of cylinders represent the relative numbers of particles. **(e)** Gold-standard Fourier Shell Correlation (FSC) curves of the final density maps. The final resolution of HKU2 spike is 2.38 Å; the final resolution of SADS-CoV spike is 2.83 Å.

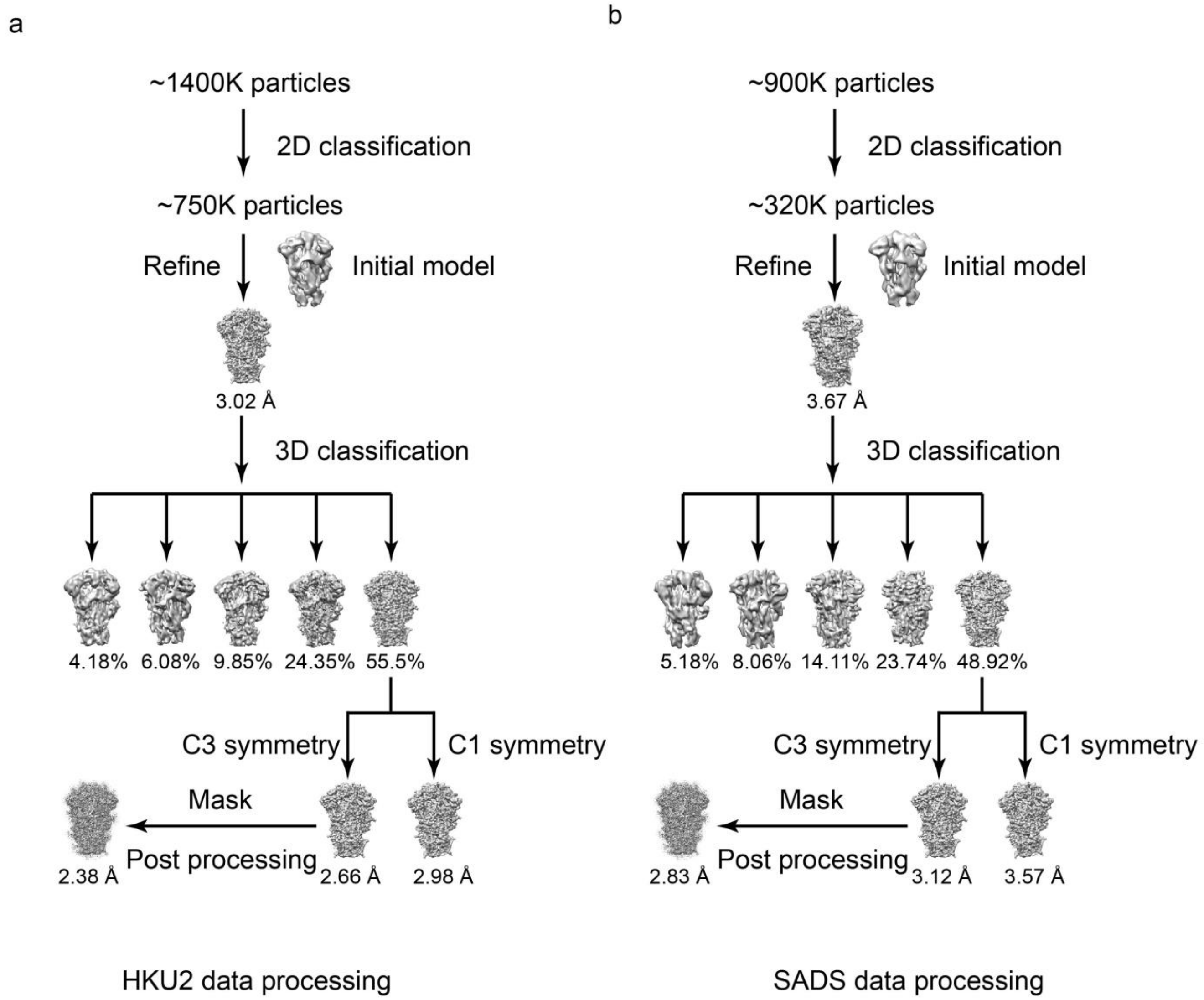


**Supplementary Fig. 2 3D reconstruction workflow. (a)** 3D reconstruction workflow of HKU2 spike cryo-EM data. **(b)** 3D reconstruction workflow of SADS-CoV spike cryo-EM data.

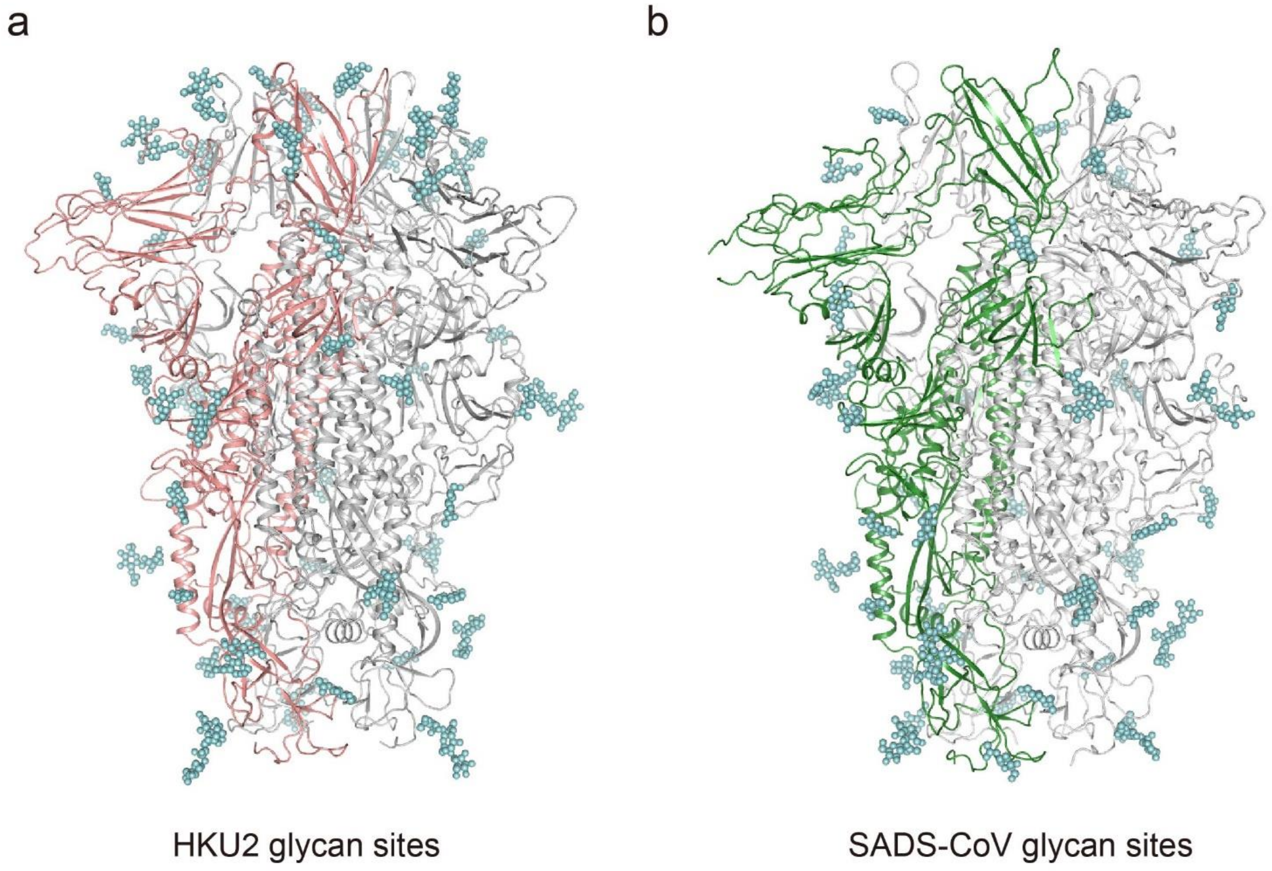


**Supplementary Fig. 3 Glycan sites of HKU2 spike and SADS-CoV spike. (a)** Glycan sites of HKU2 spike. Glycans are colored cyan. One monomer of HKU2 spike is colored salmon. The others are colored gray. **(b)** Glycan sites of SADS-CoV spike. Glycans are colored cyan. One monomer of SADS-CoV spike is colored green. The others are colored gray.

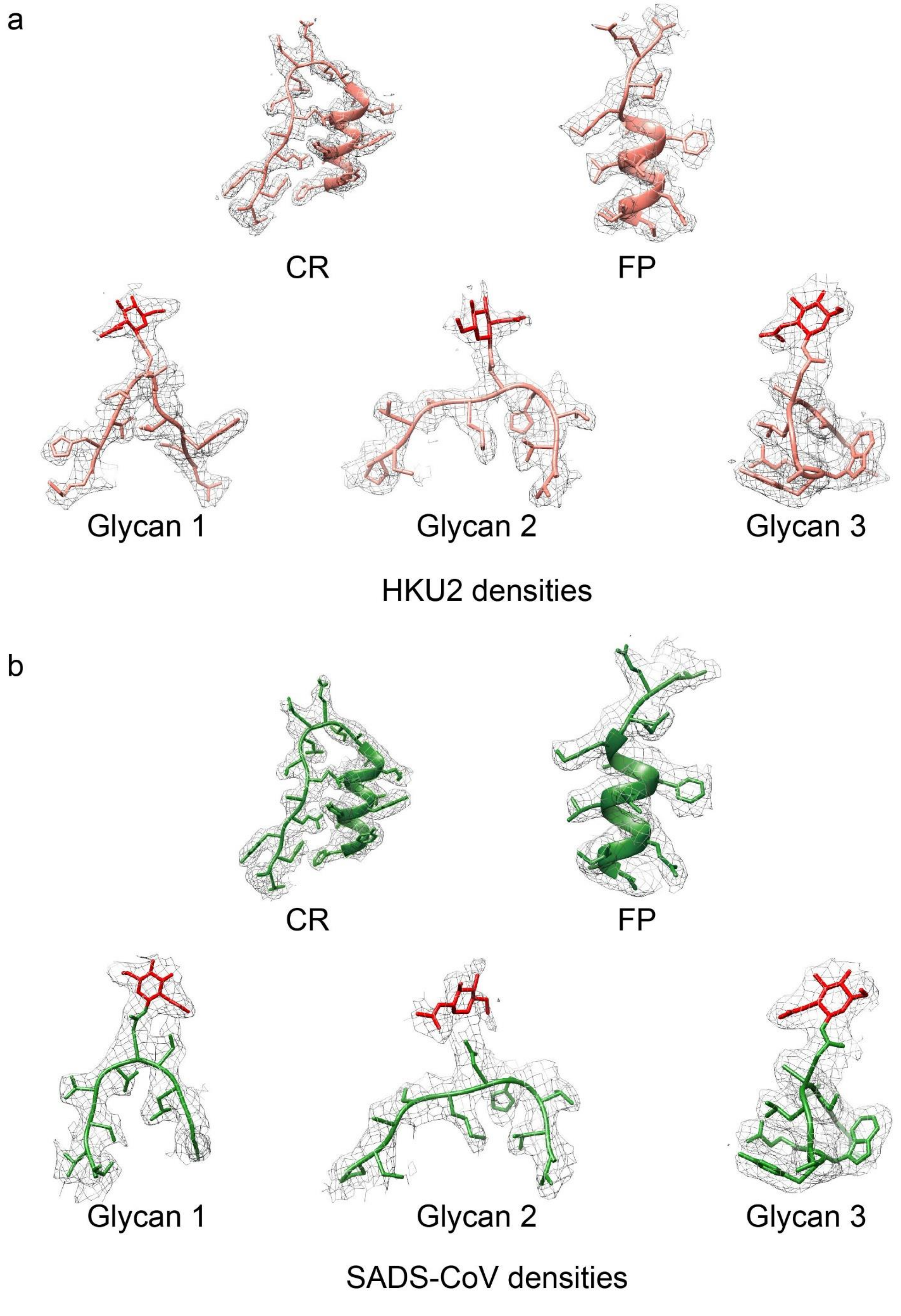


**Supplementary Fig. 4 Representative density maps. (a)** Representative density maps of HKU2 spike. The map is contoured at 2.6 RMS to show the density. CR, connecting region. FP, fusion peptide. Glycan 1 is from HKU2 NTD; glycan 2 is from HKU2 CTD; glycan 3 is from HKU2 S2. **(b)** Representative density maps of SADS-CoV spike. The map is contoured at 3

RMS to show the density. CR, connecting region. FP, fusion peptide. Glycan 1 is from SADS-CoV NTD; glycan 2 is from SADS-CoV CTD; glycan 3 is from SADS-CoV S2.

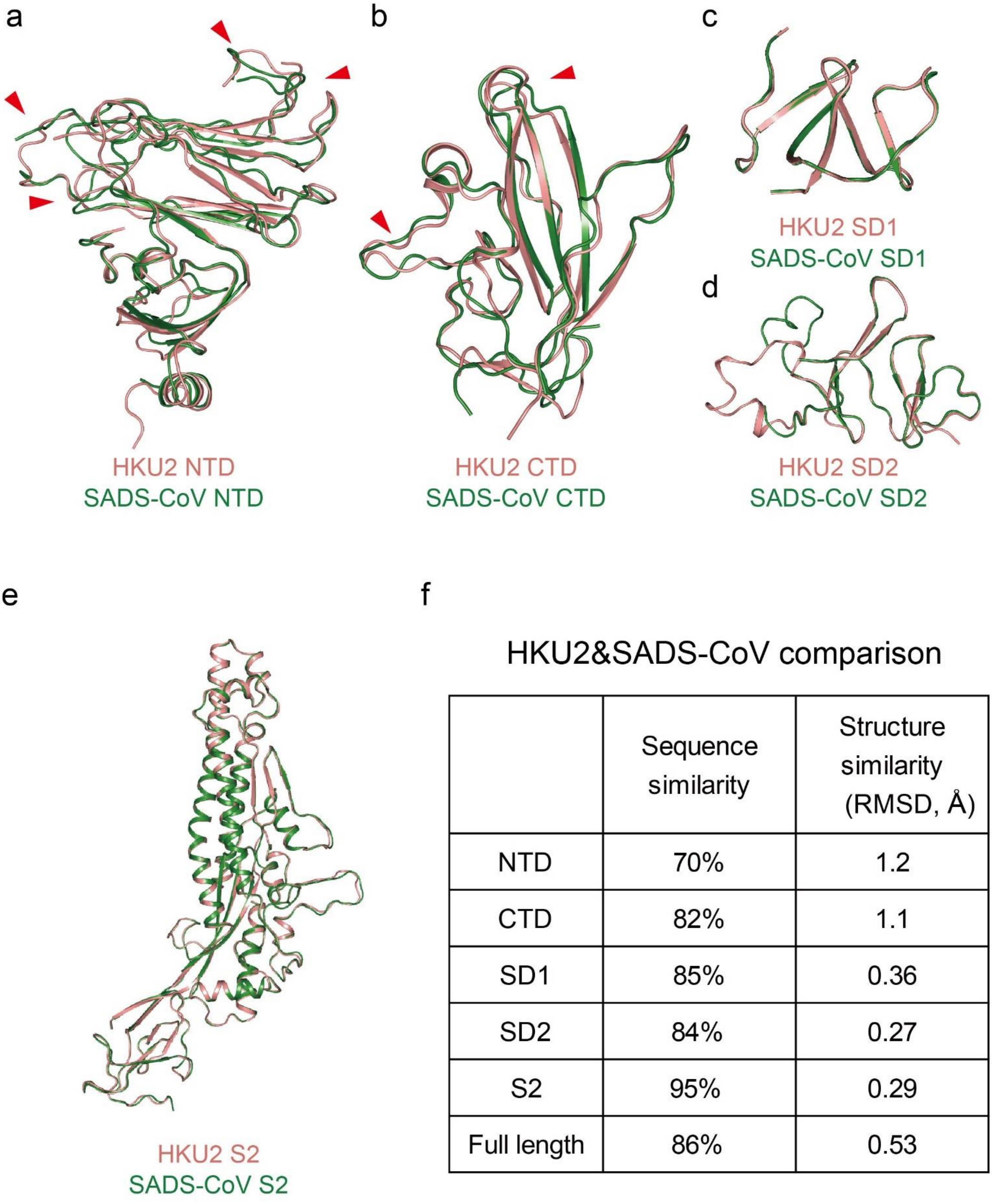


f

HKU2&SADS-CoV comparison

| | Sequence similarity | Structure similarity (RMSD, Å) |
|---|---|---|
| NTD | 70% | 1.2 |
| CTD | 82% | 1.1 |
| SD1 | 85% | 0.36 |
| SD2 | 84% | 0.27 |
| S2 | 95% | 0.29 |
| Full length | 86% | 0.53 |

**Supplementary Fig. 5 Structural comparison of HKU2 monomer and SADS-CoV monomer. (a)** Structural comparison of HKU2 NTD and SADS-CoV NTD. The variations of loops are indicated by red triangles. HKU2 NTD is colored salmon; SADS-CoV NTD is colored green. **(b)** Structural comparison of HKU2 CTD and SADS-CoV CTD. The variations of loops are indicated by red triangles. The colors are the same as in a. **(c)** Structural comparison of HKU2 SD1 and SADS-CoV SD1. The colors are the same as in a. **(d)** Structural comparison of HKU2 SD2 and SADS-CoV SD2. The colors are the same as in a. **(e)** Structural comparison of HKU2 S2 and SADS-CoV S2. The colors are the same as in a. **(f)** Quantitative comparison between segments of HKU2 spike and SADS-CoV spike. The RMSD value is calculated using PyMOL.

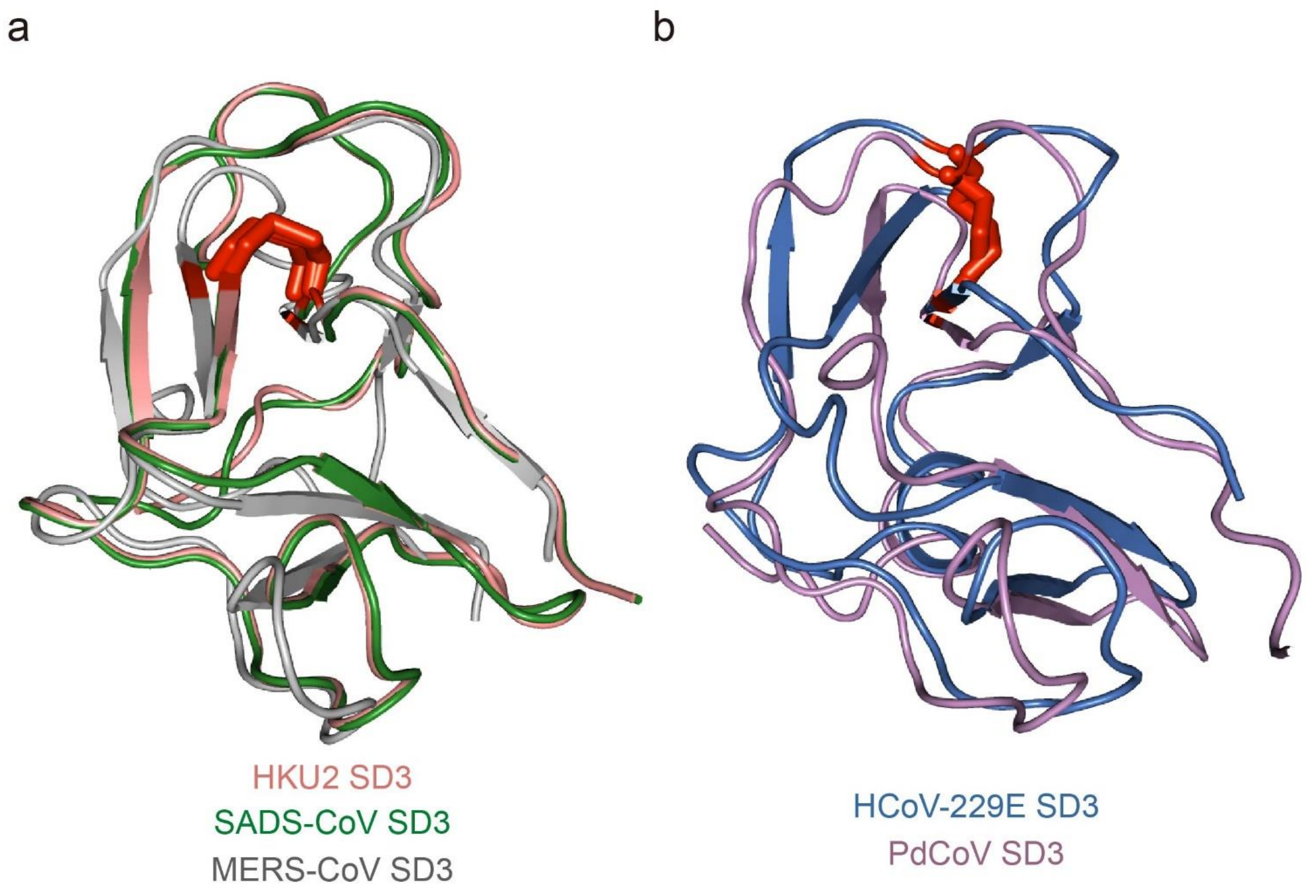


**Supplementary Fig. 6 Two types of disulfide bonds in SD3. (a)** The disulfide bond position in SD3 is the same in HKU2, SADS-CoV and MERS-CoV spikes. HKU2 SD3 is colored salmon; SADS-CoV SD3 is colored green; MERS-CoV SD3 is colored gray. PDB code: MERS-CoV, 6Q05. **(b)** The disulfide bond position in SD3 is the same in other coronavirus spikes, which are represented by HCoV-229E and PdCoV. HCoV-229E SD3 is colored marine; PdCoV SD3 is colored violet. PDB codes: HCoV-229E, 6U7H; PdCoV, 6B7N.

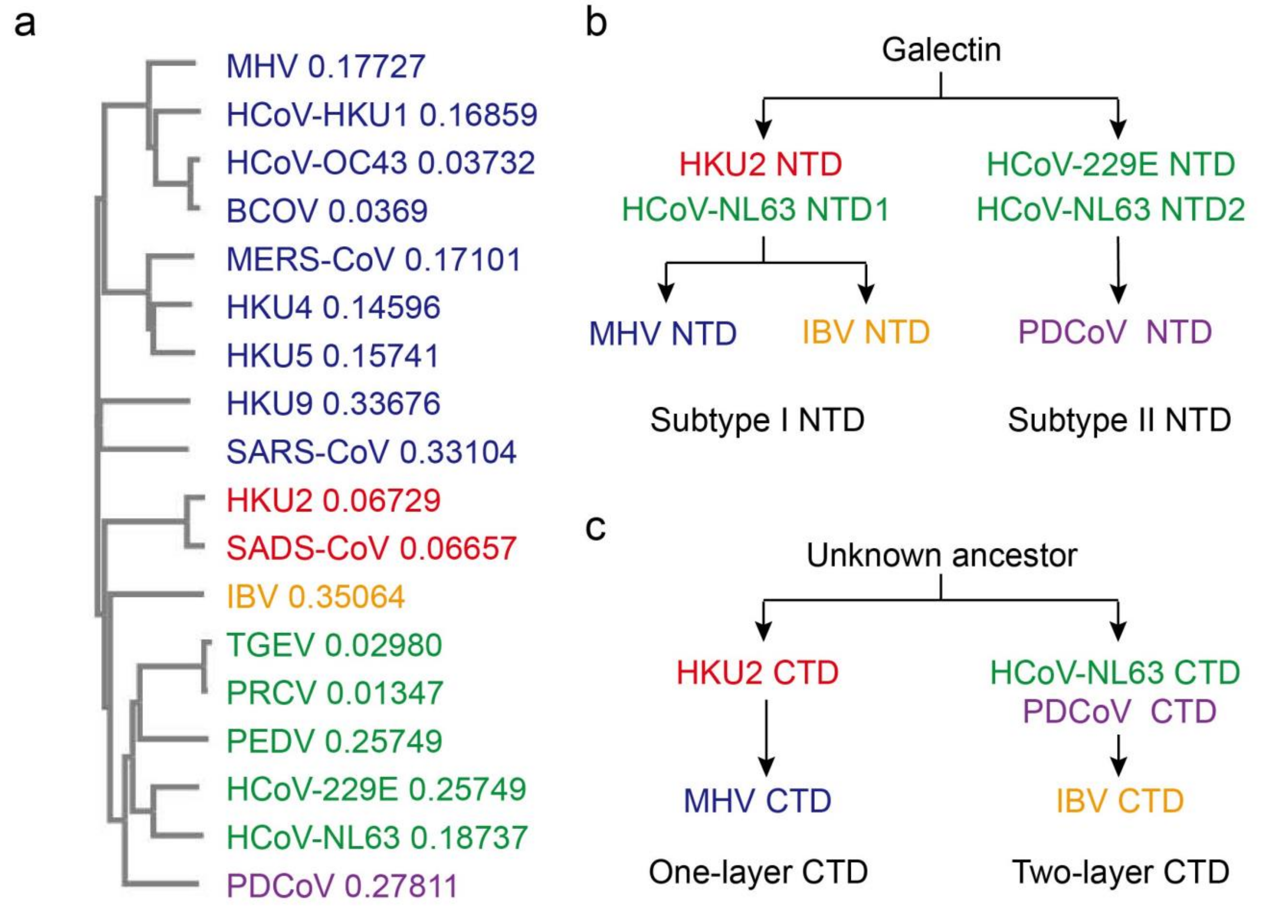


d

| RMSD(Å) | HCoV-229E | HCoV-NL63 | PEDV | TGEV | PRCV | BCoV | HCoV-HKU1 | MHV | MERS-CoV | SARS-CoV | HCoV-OC43 | IBV | PDCoV |
|---|---|---|---|---|---|---|---|---|---|---|---|---|---|
| HKU2-NTD | 4 | 2.7(I) 4.3(II) | 2.3(I) 4.1(II) | NA | NA | 3 | 3 | 2.9 | 3.2 | 3.1 | 2.8 | 3.6 | 3.9 |
| SADS-NTD | 4 | 3.1(I) 4.1(II) | 2.6(I) 4.1(II) | NA | NA | 3 | 3 | 2.9 | 3.5 | 3.5 | 3.3 | 3.5 | 3.8 |
| HKU2-CTD | 4.1 | 3.9 | 3.9 | 4.7 | 4.6 | NA | 3.4 | 3 | 3 | 3.4 | 3 | 3.5 | 4.1 |
| SADS-CTD | 4 | 3.6 | 3.5 | 4.4 | 3.9 | NA | 3.1 | 3.2 | 3.1 | 3.6 | 3.2 | 3.6 | 3.7 |
| HKU2-SD1 | 1.7 | 2 | 1.8 | NA | NA | NA | 1.7 | 2.1 | 1.9 | 1.7 | 1.6 | 1.7 | 2 |
| SADS-SD1 | 1.8 | 2.2 | 2 | NA | NA | NA | 1.7 | 2.1 | 1.9 | 1.7 | 1.6 | 1.8 | 2.1 |
| HKU2-SD2 | 2.8 | 2.4 | 2.8 | NA | NA | NA | 2.3 | 3.1 | 2.1 | 3.1 | 2 | 2.1 | 2.9 |
| SADS-SD2 | 2.8 | 2.7 | 2.9 | NA | NA | NA | 2.4 | 2.4 | 2 | 2.3 | 2.2 | 2.5 | 2.7 |
| HKU2-S2 | 4.6 | 3.5 | 3.6 | NA | NA | NA | 3 | 2.9 | 2.6 | 3.4 | 3.5 | 3.7 | 3.6 |
| SADS-S2 | 4.6 | 3.5 | 3.7 | NA | NA | NA | 3 | 2.9 | 2.7 | 3.5 | 3.5 | 3.7 | 3.7 |

**Supplementary Fig. 7 Evolution relationship of coronaviruses. (a)** Phylogenetic tree of coronaviruses. βCoVs are colored blue. HKU2 and SADS-CoV are colored red. γCoV (IBV) is colored orange. αCoVs are colored green. δCoV (PdCoV) is colored violet. The relative distances are calculated on Clustal Omega. **(b)** Putative evolution pathways of coronavirus NTDs. The colors are the same as in a. **(c)** Putative evolution pathways of coronavirus CTDs. The colors are the same as in a. **(d)**

Quantative comparison between segments of HKU2 (SADS-CoV) and coronaviruses from different genera. The colors are the same as in a. The RMSD values are calculated on DALI server.

#### Supplementary Table 1 Data Collection and Refinement Statistics

| Parameter | HKU2 Spike | SADS Spike |
|---|---|---|
| **Data collection** | | |
| Microscope | Titan Krios | Titan Krios |
| Voltage (kV) | 300 | 300 |
| Defocus range(um) | 1-3 | 1-3 |
| No. of movies | 7663 | 4568 |
| Frames per movie | 32 | 32 |
| Exposure time per frame(ms) | 175 | 175 |
| Magnification | 130000× | 130000× |
| Dose rate ($e^{-}/Å^{2}/s$) | 8.89 | 8.89 |
| Total dose per movie ($e^{-}/Å^{2}$) | 49.784 | 49.784 |
| **Data processing** | | |
| No. of particles | 421490 | 152334 |
| Symmetry | C3 | C3 |
| Provided B factor (Å2) | -87.05 | -101.51 |
| Map resolution (Å) | 2.38 | 2.83 |
| **Model validation** | | |
| Correlation coefficient(CCmask) | 0.88 | 0.86 |
| EMRinger score | 5.62 | 3.91 |
| MolProbity score | 1.77 | 2.38 |
| All-atom clash score | 4.03 | 7.8 |
| Rotamers outliers (%) | 3.58 | 3.6 |
| Ramachandran favored (%) | 97.08 | 90.55 |
| Ramachandran allowed (%) | 2.92 | 9.45 |
| Ramachandran outliers (%) | 0 | 0 |
| **$\mathbf{RMSD}^{a}$** | | |
| Bond length (Å) | 0.008 | 0.009 |
| Bond angles (°) | 0.737 | 0.783 |

$\mathbf{RMSD}^{a}$, root mean square deviations.
